# Chromatin accessibility of primary cancers informs regional mutagenesis in metastases through multi-scale deep learning

**DOI:** 10.64898/2026.05.24.727503

**Authors:** Hanli Jiang, Alexander T. Bahcheli, Kevin C.L. Cheng, Jüri Reimand

## Abstract

Tissue-specific chromatin states shape regional mutation density in primary tumors, but whether this relationship persists in metastases is unclear. We integrated whole-genome sequencing data from 2,507 metastatic tumors across six cancer types with 892 chromatin accessibility and replication timing profiles, and developed CAMM, a multi-scale deep learning model, to jointly predict SNV and indel density across multiple genomic resolutions. The model explained regional variance in mutation density across metastatic cancers and preserved mutagenesis patterns in an independent primary tumor cohort. Chromatin accessibility profiles from tissue-matched primary cancers were informative predictors of regional mutagenesis in metastases, supporting a contribution of lineage-linked chromatin context, as indicated by feature attribution analyses. Genomic windows with mutation burdens exceeding epigenome-based expectations were enriched for known and candidate cancer loci. These results link metastatic regional mutagenesis to tissue-of-origin chromatin accessibility and provide a framework for interpreting mutational processes and prioritizing mutation-enriched loci.

## Introduction

Metastasis is the major cause of cancer mortality and remains the central barrier to durable cure ^1–3^. Although metastatic lesions arise from a primary tumor, they often evolve under new selective pressures during dissemination and colonization and can acquire additional mutations associated with progression and therapy exposure ^1,4^. An open question is whether the mutational processes that shape metastatic genomes still reflect the chromatin context of the tissue in which the cancer originated, or whether these patterns are substantially remodeled during advanced disease. Resolving this question matters for two reasons: it helps to distinguish which features of the metastatic mutation landscape are retained from the tissue of origin versus newly acquired, and it improves our ability to interpret mutation enrichment as evidence of selection rather than background mutagenesis.

Somatic mutations, including single-nucleotide variants (SNVs) and short insertions-deletions (indels) shape the cancer genome and reflect its evolutionary history and molecular environment ^5,6^. A small subset of these alterations function as drivers that confer a selective advantage ^7–9^; however, the majority of mutations are passenger events produced by diverse mutational processes ^10–12^. Accurately characterizing mutational processes is therefore essential for reconstructing tumor evolution and for distinguishing signals of selection from variation in background mutation rates ^13,14^. Mutational heterogeneity is evident across multiple genomic scales. At base-pair resolution, single-base substitution signatures reflect processes such as aging, exogenous mutagens, defects in DNA repair, and treatment-associated damage, each with characteristic sequence-context preferences ^10,15^. At intermediate scales, specific regulatory elements such as transcription start sites and CTCF binding sites are subject to distinct mutational processes that can reflect local DNA breakage and repair activity ^16–18^. At kilobase-to-megabase scales, regional mutation density varies widely and correlates with functional genomic features including replication timing (RT), chromatin accessibility (CA), and transcriptional activity ^19–23^. Early-replicating, transcriptionally active regions of open chromatin show fewer mutations than late-replicating and closed regions, possibly due to differential error rates and mismatch repair activity during DNA replication ^20,24–27^. Importantly, because chromatin organization is strongly tissue-specific, CA profiles measured in primary tumors are highly predictive of regional mutation density across many cancer types ^22,23,28^. This observation implies that regional mutagenesis can carry information about the tumor cell-of-origin and that interpretable predictive models can reveal which epigenomic contexts best explain mutation rate variation ^23^.

Whether these epigenomic-to-mutagenic relationships persist in metastatic tumors is not fully understood. Metastases can experience additional selective pressures during dissemination and outgrowth, and treatment can introduce strong mutational footprints that differ from endogenous processes ^4,15^. These factors could weaken chromatin-linked patterns learned from primary tumors. At the same time, many features of chromatin organization are maintained across cell divisions and may continue to constrain mutation accumulation even after dissemination. Despite this, there has been no systematic evaluation of regional mutagenesis in metastatic tumors using an interpretable and consistent modeling framework.

Here, we asked whether tissue-of-origin epigenomic features continue to shape regional mutagenesis in metastatic cancer and whether excess mutation burden relative to epigenome-based expectation can help identify mutation-enriched loci. We integrated whole-genome sequencing data from metastatic tumors of multiple cancer types with hundreds of chromatin accessibility and replication timing profiles. To capture regional mutagenesis across genomic scales, we developed the Chromatin Accessibility-informed Multi-scale Mutagenesis Model (CAMM), a deep learning framework that predicts regional SNV and indel density from megabase to kilobase windows. We leveraged this framework to study how tissue-of-origin chromatin accessibility relates to regional mutagenesis and to identify coding genes and non-coding elements with excess mutation burden. These analyses support a persistent influence of tissue-of-origin chromatin context on the mutational landscape of metastatic cancer.

## Results

### Epigenomic landscapes capture regional mutation density across metastatic genomes

We first asked whether epigenomic features that explain regional mutation density in primary tumors also predict regional mutation density in metastases. To this end, we analyzed whole-genome sequencing (WGS) data from 2,507 metastatic tumors across six cancer types (breast, colorectal, prostate, lung, esophagus, and skin) in the Hartwig Medical Foundation (HMF) cohort ^4^, comprising 45.4 million SNVs and 3.5 million indels in total (**Table S1**). We combined these data with CA profiles from 796 primary tumor profiles in the TCGA ATAC-seq dataset ^29^ and 96 RT profiles representing various common cell lines from ENCODE Repli-seq experiments ^30^ (**Table S2**).

To model regional mutagenesis, we summarized SNVs and indels in non-overlapping genomic windows at 1 Mb, 100 kb, and 10 kb resolutions. For each window, we computed the corresponding CA and RT signal values from each epigenomic profile, yielding a window-by-feature matrix that links regional epigenomic context to regional mutation density (**Fig. 1A**). This window-level formulation allowed us to ask how well regional epigenomic context predicts mutation density in metastatic cancer genomes.

**Figure 1.**
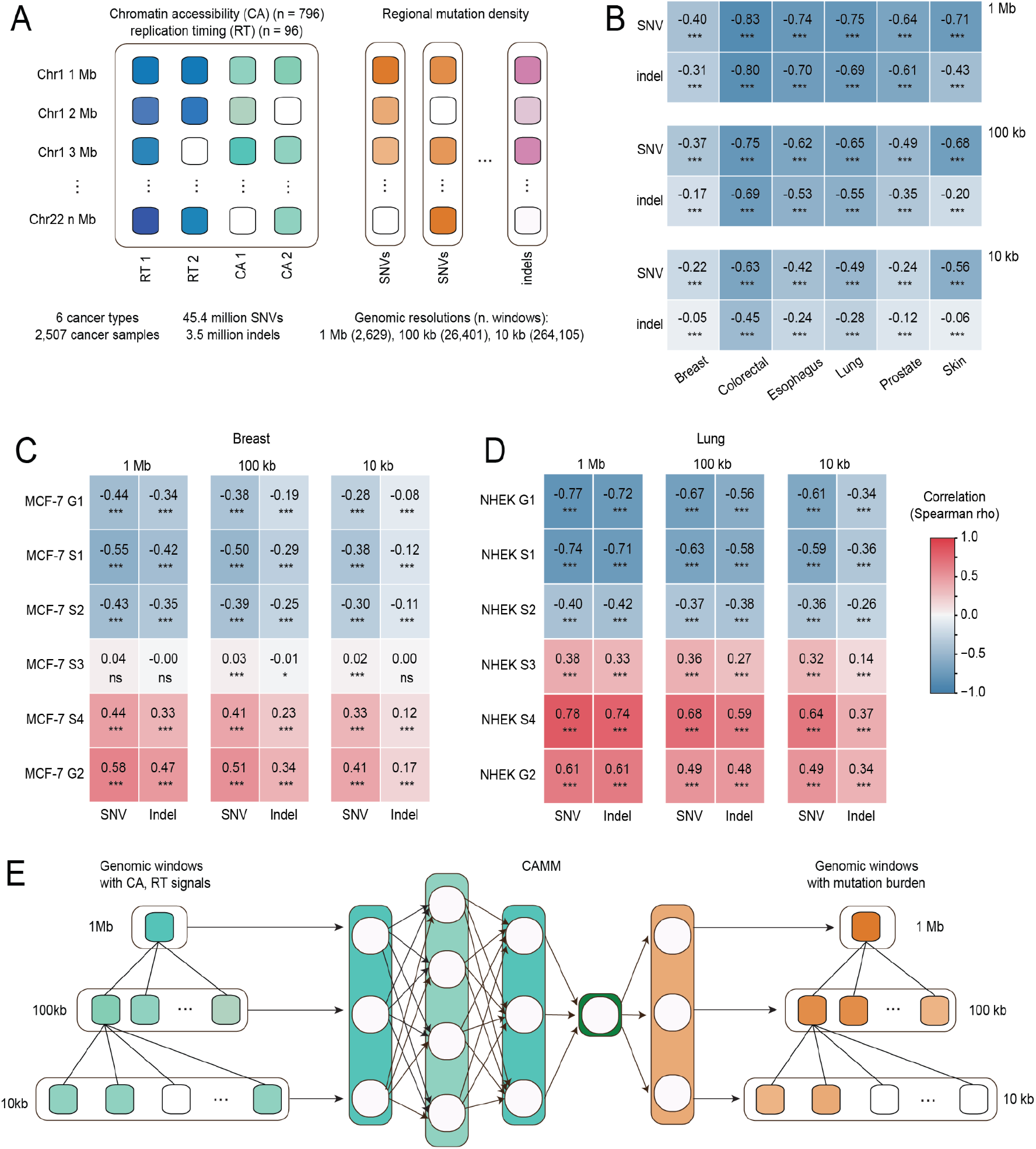
Overview of data and model design. **(A)** Window-level representation of epigenomic inputs and mutation outputs. Mean chromatin accessibility (CA; green) and replication timing (RT; blue) signals were computed for genomic windows at 1 Mb, 100 kb, and 10 kb resolutions. Regional mutation density was defined as the number of SNVs or indels per window, generating paired input-output matrices for supervised learning across six cancer types. **(B)** Spearman correlations between CA profiles and regional SNV or indel density matched by cancer type. CA was consistently negatively correlated with mutation density across cancer types and resolutions. Colors indicate Spearman’s ρ. **(C and D)** Spearman correlations between cancer-type-matched RT state and mutation density in breast and lung cancer contexts. Early-replicating regions correlated negatively with mutation density, whereas late-replicating regions correlated positively, consistent across SNVs and indels and across multiple window resolutions. Asterisks indicate P-values: *** P < 0.001, ** P < 0.01, * P < 0.05, and ns, not significant. **(E)** CAMM model architecture. CA and RT features from 1 Mb, 100 kb, and 10 kb genomic windows were integrated in a hierarchical multi-scale, multi-task framework. Scale-specific embeddings were learned at each resolution, coarser-scale information was propagated to finer scales, and SNV and indel burden were jointly predicted across resolutions.

Before model training, we examined the marginal relationship between epigenomic features and mutation density. Cancer-type-matched CA profiles were consistently negatively correlated with regional SNV and indel density across cancer types and genomic resolutions, indicating that more accessible chromatin regions generally accumulated fewer mutations, extending observations from primary cancers ^22,23^ (**Fig. 1B**). RT profiles showed the expected directional relationship with mutation density. In representative breast and lung cancer contexts, early-replicating states were negatively correlated with SNV and indel density across multiple genomic resolutions whereas late-replicating states were positively correlated, consistent with prior observations linking RT to mutation density in primary cancers ^21,26,27^ (**Fig. 1C-D**). Together, these correlation analyses confirmed that both CA and RT retain meaningful associations with regional mutation density in metastatic cancer and motivated their use as predictive features.

### Multi-scale, multi-task learning using CAMM improves mutation density prediction

Regional mutation density is shaped by constraints acting across multiple genomic scales, from broad chromatin domains to local regulatory elements ^13,14,20,23^. To capture this nested structure, we implemented CAMM, a hierarchical multi-scale deep learning model that propagates information from coarse (1 Mb) to intermediate (100 kb) to fine (10 kb) genomic resolutions (**Fig. 1E**). At each scale, CAMM first learns scale-specific representations of CA and RT features and then passes coarser-scale information forward to finer scales, allowing local predictions to be informed by broader genomic context. This coarse to fine information flow was inspired by U-Net-like architectures, which combine broad contextual representations with finer-resolution features through hierarchical information transfer. We adapted this structure from image segmentation to genomic-window prediction by passing coarser epigenomics information forward to finer-scale mutation prediction layers ^31^. In parallel, because SNVs and indels share several genomic correlates including CA and RT ^20^, we used a multi-task formulation that jointly predicts SNV and indel density while allowing task-specific behavior at fine scale (**Methods**) ^19^. This architecture enables CAMM to integrate broad and local epigenomic signals while leveraging shared structure between mutation classes, making it well suited to model regional mutagenesis across both genomic scales and mutation types.

To evaluate model performance, we first quantified the contribution of multi-scale context by comparing multi-scale models to single-scale models trained independently at each resolution. Across cancer types, the multi-scale design improved SNV prediction at all resolutions, with the largest gains at 100 kb resolution (**Fig. 2A**), consistent with this scale balancing domain-level context and sufficient mutational signal per window. For indels, multi-scale context yielded smaller gains, particularly at 10 kb (**Fig. S1B**), consistent with indel density being more sensitive to fine-scale determinants (including local sequence features) that are not directly represented by regional CA/RT summaries ^10^. Notably, for SNVs, a combined multi-scale model integrating information across all resolutions improved prediction beyond models trained at any individual resolution alone, indicating that complementary predictive signals are captured across scales.

**Figure 2.**
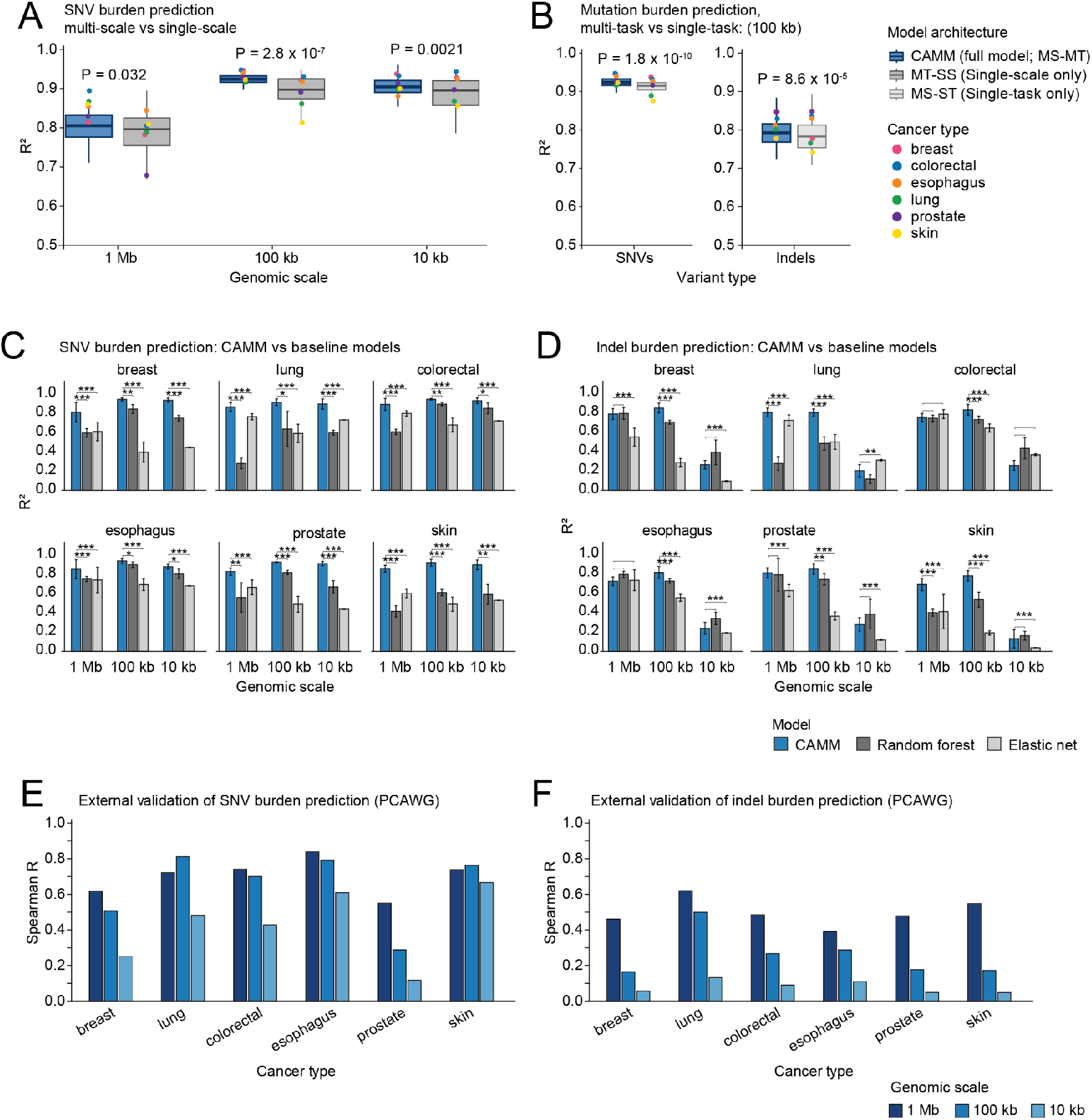
CAMM improves prediction of regional mutation density and transfers to independent primary cancer data. **(A)** Comparison of SNV prediction performance in HMF between CAMM, the full multi-scale, multi-task model (MS-MT), and the single-scale baseline (MT-SS) across three genomic window sizes (1 Mb, 100 kb, and 10 kb). Boxplots summarize R^2^ values across cancer types, and colored points indicate the median R^2^ for each cancer type. **(B)** Comparison of CAMM (MS-MT) with the single-task baseline (MS-ST) at 100 kb resolution for SNVs (left) and indels (right), showing the effect of joint learning across mutation types. **(C-D)** Prediction performance of CAMM, random forest, and elastic net models for **(C)** SNVs and **(D)** indels across 1 Mb, 100 kb, and 10 kb windows. Bars show mean R^2^ across cancer types, and error bars indicate variability across runs. Asterisks indicate significance annotations for pairwise comparisons between CAMM and baseline models (*** P < 0.001, ** P < 0.01, * P < 0.05). **(E-F)** External validation in PCAWG primary tumors showing Spearman correlation between predicted and observed regional mutation density across cancer types at 1 Mb, 100 kb, and 10 kb resolutions for **(E)** SNVs and **(F)** indels. Bars show correlations for individual cancer types, with positive correlations indicating preservation of the rank ordering of regional mutation density in an independent cohort of primary tumors.

We next evaluated the benefit of joint SNV-indel learning by comparing multi-task models and single-task models (**Fig. S1C-D**). Multi-task learning improved both SNV and indel prediction, again most clearly at 100 kb where SNV and indel patterns co-vary strongly across windows (**Fig. 2B**). Together, these comparisons support the two core modeling choices underlying downstream biological interpretation: integrating multi-scale genomic context and sharing information across mutation types.

### Accurate prediction of regional mutation density across metastatic cancer types

CAMM achieved strong predictive performance across metastatic cancer types by combining multi-scale and multi-task learning (**Fig. 2C-D**). For SNVs, prediction accuracy was high at all three resolutions, with R^2^ of 0.80-0.90 at 1 Mb, 0.90-0.93 at 100 kb, and 0.80-0.90 at 10 kb (**Fig. 2C**). To benchmark CAMM against simpler alternatives, we compared it with both a non-linear tree-based model (random forest) and a regularized linear model (elastic net). At each resolution, CAMM outperformed both baselines, indicating that the learned non-linear integration of CA and RT features improves prediction of regional mutation density beyond both standard tree-based and regularized linear models (**Fig. 2C**). For indels, prediction accuracy was lower than for SNVs, particularly at 10 kb resolution (R^2^ of 0.69-0.80 at 1 Mb, 0.79-0.85 at 100 kb, and 0.12-0.28 at 10 kb; **Fig. 2D**). This reduced performance at 10 kb is not unexpected and is consistent with two constraints: first, indel counts are substantially sparser than SNVs at 10 kb resolution, and second, fine-scale indel variability is influenced by sequence-level properties that are not optimally captured by regional CA and RT features ^10,11^. Despite this limitation, indel prediction remained consistently high-quality at 1 Mb and 100 kb, supporting a detectable contribution of chromatin context to indel density at broader scales.

As a stricter test of genomic generalization, we performed chromosome-held-out validation by training CAMM on 21 autosomes and testing on the remaining held-out chromosome. This process was repeated across all autosomes, cancer types, mutation classes, and resolutions. Most held-out chromosomes retained strong predictive performance, indicating that CAMM was not solely relying on correlations among neighboring windows from random splits (**Fig. S2 A-B**). Some cancer types showed reduced or negative R^2^ on one or two chromosomes because outlier windows affected absolute calibration, but Spearman correlations remained consistently positive, supporting preserved rank ordering of regional mutation density. In prostate cancer, chromosome 8 showed outlier-driven R^2^ loss, consistent with the frequent copy-number alteration of chromosome 8 in advanced prostate cancer, including recurrent 8q gain involving *MYC* and 8p loss involving *NKX3-1* ^32,33^. Chromosome 14, by contrast, showed strong and robust performance (**Fig. S2 C-D**).

Together, these results show that CAMM provides strong predictive performance for regional mutation density, especially for SNVs and broader-scale indel prediction, while outperforming simpler linear and tree-based baselines.

### Epigenome-based mutagenesis models transfer from metastases to primary tumors

To test whether CAMM trained on metastatic genomes captures broader features of regional mutagenesis beyond cohort-specific effects, we evaluated transfer to primary, untreated tumors from the Pan-cancer Analysis of Whole Genomes (PCAWG) cohort ^5^. For consistency with the metastatic analysis, we focused on the same six cancer types (breast, colorectal, prostate, lung, esophagus, and skin), comprising 889 primary tumors with 9.3 million SNVs and 0.45 million indels (**Table S3**). Because absolute mutation burden differs between metastatic and primary cohorts, we assessed transfer using non-parametric Spearman correlation between observed and predicted regional mutation density.

CAMM preserved the rank ordering of genomic windows by mutation density in matched PCAWG primary tumor cohorts even without retraining. At 1 Mb resolution, the model achieved median Spearman correlations of ρ = 0.72 for SNVs and ρ = 0.50 for indels (**Fig. 2E-F**). When genomic windows were pooled across cancer types, both correlations remained highly significant (Spearman P < 2.2 x 10^-16^). Transfer performance was lower in PCAWG than in HMF, likely owing to differences in mutation burden, cohort composition, data processing, and disease context, including treatment exposure ^4,5,28,34^. Despite these differences, positive correlations across cancer types and scales indicate that the epigenome-mutagenesis relationships learned in metastases are not restricted to the metastatic setting.

### Tissue-matched chromatin accessibility profiles are enriched among top predictors in most metastatic cancer types

We next asked which epigenomic profiles contribute most strongly to mutation-density prediction in metastases. We examined feature importance in CAMM, which jointly integrates predictors across multiple genomic scales to capture both broad and local determinants of regional mutation density (**Fig. 3**). Using SHAP-based feature attribution ^35^ and custom permutation-based importance testing, we ranked CA and RT profiles by their contribution to overall model performance and individual regional predictions (**Methods**).

**Figure 3.**
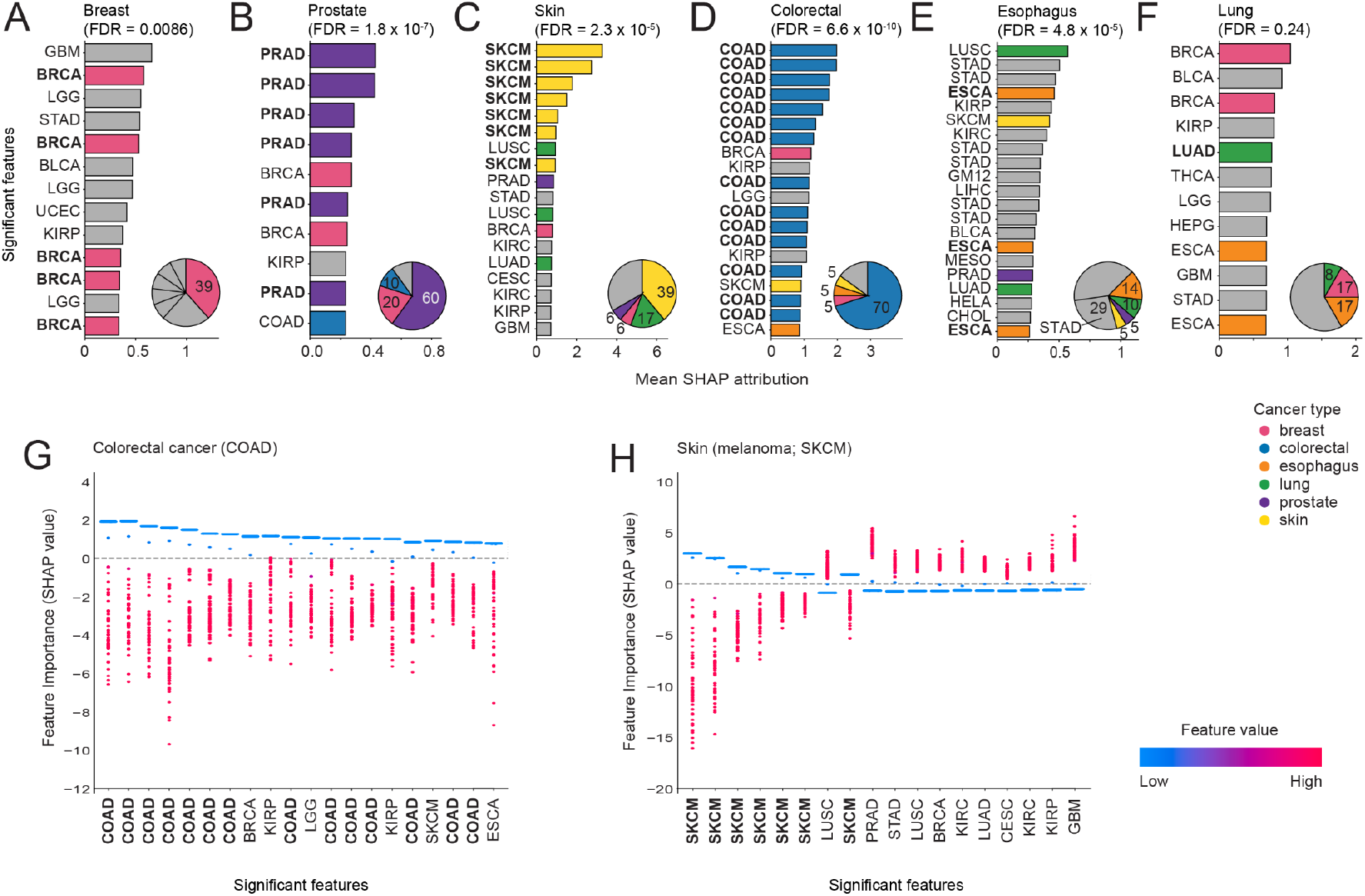
Tissue-matched chromatin accessibility profiles are the dominant predictors of regional mutation density in metastatic cancers. **(A-F)** Ranked significant epigenomic features for each metastatic cancer type, ordered by mean absolute SHAP attribution from the CAMM. Bars are colored by the tissue or cancer type represented by each feature. Inset pie charts show the composition of top features across cancer type groups, with percentages of relevant CA profiles shown in the pie charts. Permutation tests were used to determine significance (P < 0.001). **(G-H)** SHAP value distributions for representative significant predictors in colorectal and skin metastases, showing the contribution of feature values across individual genomic regions. Negative SHAP scores on the Y-axis show reduced mutation burden associated with corresponding CA features while positive scores reflect positive associations with mutation burden.

Across cancer types, CA profiles were more often selected as significant predictors and showed larger attributions than RT profiles (**Fig. 3A-F**). This likely reflects the distinct biological properties of these features: RT is relatively conserved across cell types and varies mainly over broad genomic domains, whereas CA captures richer tissue-specific regulatory information and stronger local variation. As a result, although the model incorporates both broad- and fine-scale genomic context, tissue-matched CA profiles remain the most informative predictors of regional mutation density in metastatic cancers. This is consistent with prior work showing that replication timing is most relevant at larger genomic scales ^36–38^, whereas chromatin accessibility better resolves local mutational variation ^22^.

We next asked whether the most informative epigenomic predictors of regional mutagenesis in metastases reflect tissue-of-origin and lineage-specific chromatin states. Extending prior observations from primary tumors ^22,23^, the most influential CA features in metastases were not generic measures of chromatin openness; instead, they were strongly enriched for tissue-matched profiles. To formally test this observation, we classified CA profiles as tissue-matched, anatomically related, or unrelated for each cancer type and compared the top predictors against the full background of 796 CA profiles. In breast, prostate, skin, and colorectal metastases, CA profiles from the corresponding primary tumor types were prominent among the top-ranked predictors (**Fig. 3A–D**). This pattern was supported by both stronger enrichment of tissue-matched profiles among top predictors and higher attribution magnitudes across the full CA profile set (FDR < 0.01 for all Fisher’s exact and Wilcoxon tests). Esophageal and lung cancer metastases showed more nuanced patterns, prompting separate interpretation of these primary cancer contexts. When matched esophageal carcinoma (ESCA) profiles were considered together with anatomically related CA profiles from upper-gastrointestinal cancers, including stomach adenocarcinoma (STAD), these matched-or-related CA profiles remained enriched among top predictors and showed higher attribution magnitudes than unrelated profiles (**Fig. 3E**; FDR < 0.01 for both tests). In contrast, lung cancer metastases showed no significant enrichment of tissue-matched or anatomically related CA profiles (**Fig. 3F**; Fisher FDR = 0.24; Wilcoxon FDR = 0.72), suggesting a broader and less lineage-restricted feature-importance landscape. Together, these results support enrichment of tissue-matched or anatomically related CA profiles among influential predictors in most metastatic cancer types.

We next asked whether tissue-matched CA profiles show consistent directional effects on regional mutagenesis across individual genomic regions. In colorectal and skin metastases, higher CA in tissue-matched profiles was associated with negative SHAP attributions, whereas lower CA showed the opposite pattern (**Fig. 3G-H, Fig. S3**). This pattern was particularly clear in skin metastases, where the tissue-matched skin cutaneous melanoma (SKCM) CA profile showed a consistently negative association, whereas unrelated CA features did not. This association between CA and regional mutation burden is consistent with previous studies in primary cancers ^20,22,23^. Together, these analyses indicate that tissue-linked CA profiles remain the most informative epigenomic predictors of regional mutation density in the metastatic setting.

### Mutation-enriched loci at known cancer genes pinpointed by model residuals

Driver mutations arise under positive selection and can appear in large cancer cohorts as excess mutation burden beyond background expectation ^7,9^. Because CAMM captures major epigenomic determinants of regional mutation density ^20,23^, its residuals provide a way to identify genomic windows with mutation burdens that exceed regional epigenomic expectation ^39^. We therefore asked whether genomic windows with high positive residuals preferentially overlap cancer-relevant genomic elements. We focused this analysis on 10-kb windows, which provide the highest locus-level resolution in our framework and enable more precise assignment of enriched windows to nearby genes and non-coding elements.

To identify a high-confidence set of mutation-enriched windows, we focused on genomic windows with especially high model residual values. Residuals were first log-transformed and then converted to standardized Z scores, and a cutoff of Z > 4 was used to define high-confidence outliers. We identified a high-confidence set of 58 mutation-enriched 10-kb windows (**Table S4**), many of which overlapped known cancer-gene loci ^40,41^. This corresponded to a six-fold enrichment over chance expectation (18/58 or 31%; Fisher’s exact P = 1.6 x 10^-10^), whereas the remaining windows highlighted additional candidate coding and non-coding loci of potential interest (**Fig. 4A-B**). Thus, genomic loci with mutation burden exceeding epigenome-based expectation are enriched for cancer-relevant elements. To define this high-confidence set, we used a stringent standardized residual Z-score filter (Z>4). To test whether this enrichment depended on the residual threshold, we repeated the analysis using different Z-score thresholds and found that lower thresholds selected more outlier windows, whereas higher thresholds retained smaller, more stringent sets (**Fig. 4C**). Enrichment for known cancer-gene loci was maintained across thresholds, supporting the robustness of the residual signal. We used the threshold Z>4 for downstream analyses because it retained a focused set of mutation-enriched windows while preserving strong cancer-gene enrichment.

**Figure 4.**
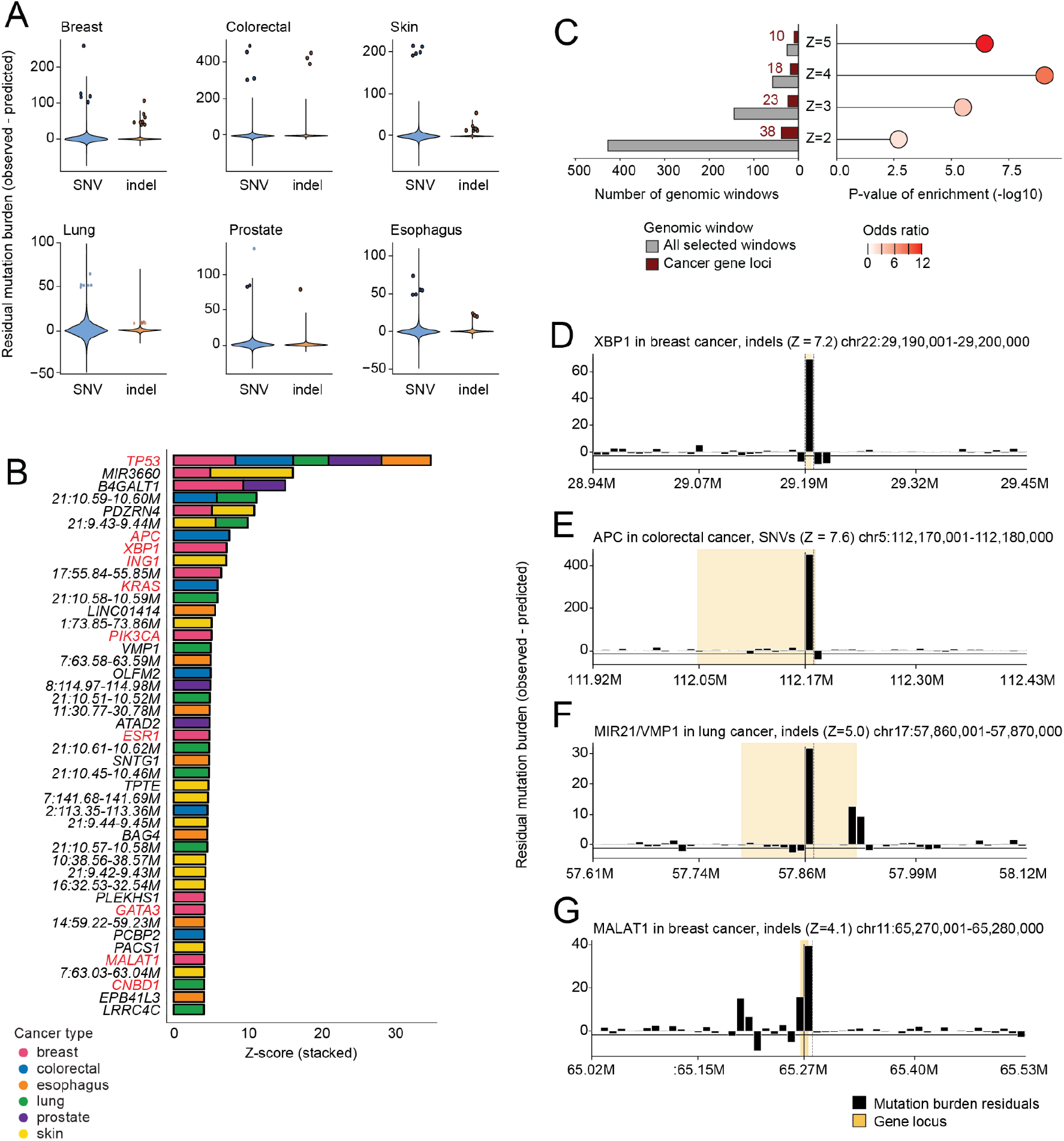
Residual-based analysis identifies mutation-enriched genomic windows at known cancer genes and candidate loci. (A) Distributions of model residuals across 10-kb genomic windows for each cancer type and mutation class. Violin plots show genome-wide residual distributions, and points indicate outlier windows, corresponding to regions where observed mutation burden exceeded epigenome-based prediction. (B) Genes and genomic regions overlapping high-residual outlier windows, ranked by summed residual Z score across cancer types and mutation classes. Outlier windows that overlap annotated genes are labeled by gene name, whereas intergenic or unannotated regions are labeled by chromosome coordinates; for example, 21:10.59–10.60M denotes a region on chromosome 21 spanning 10,590,000–10,600,000 bp. Bars are stacked by cancer type, and known cancer genes are highlighted in red. (C) Sensitivity analysis of residual Z-score thresholds used to define mutation-enriched outlier windows. The left panel shows the number of selected windows and the subset overlapping known cancer-gene loci across thresholds from Z > 2 to Z > 5. The right panel shows enrichment of cancer-gene overlap at each threshold, with the x-axis showing −log10-transformed Fisher’s exact test P values and point color indicating the odds ratio. The Z > 4 threshold was used for downstream analyses to prioritize high-confidence mutation-enriched windows while retaining strong enrichment for known cancer genes. (D–G) Representative mutation-enriched loci identified by residual analysis. Bars show residuals for adjacent 10-kb windows, calculated as observed minus predicted mutation burden. Dashed vertical lines mark the focal high-residual window, and shaded regions indicate annotated gene loci where shown. Examples include XBP1 in breast cancer indels (D), APC in colorectal cancer SNVs (E), the VMP1 region in lung cancer indels (F), and the MALAT1/TALAM1/MASCRNA non-coding RNA locus in breast cancer indels (G).

Examining the top mutation-enriched loci, *TP53* (**Fig. S4**) was the most recurrently implicated gene with mutation enrichment in five cancer types (**Fig. 4A-B**). Several additional loci showed tumor-type-specific enrichment patterns that recapitulated established cancer biology. In breast metastases, *ESR1* and *PIK3CA* showed strong mutational enrichment (**Fig. 4B**), consistent with their known roles in disease progression and therapy resistance ^42,43^. Breast metastases also highlighted *XBP1* (**Fig. 4D**), a key transcription factor in the unfolded protein response that has been implicated in triple-negative breast cancer progression ^44^. In colorectal metastases, prominent mutational enrichment occurred at *APC* (**Fig. 4E**) as well as *KRAS*, consistent with canonical colorectal cancer genetics ^45,46^. In skin cancer metastases, *ING1* showed marked mutational enrichment, consistent with its role as a tumor suppressor linked to UV damage responses and nucleotide excision repair ^47^. As a candidate example, *ATAD2* emerged as a mutation-enriched locus in prostate metastases, consistent with its role as a MYC-associated chromatin regulator linked to aggressive tumor behavior ^48,49^.

Beyond well-known cancer genes, residual outlier windows also highlighted candidate non-coding loci. One such locus encompassed *VMP1* (**Fig. 4F**) and *MIR21*, including the well-established oncogenic microRNA *MIR21* ^50^, Prior work has also described regulatory coupling between miR-21 and VMP1 in colorectal cancer cells ^51^. We also identified the non-coding RNA locus *MALAT1* (**Fig. 4G**) in breast cancer, consistent with a metastasis-related role in breast cancer and with recurrent non-coding mutation enrichment in cancer genomes ^8,52^. Together, these findings suggest that residual analysis complements epigenome-based prediction by prioritizing loci whose mutation burden is not fully explained by chromatin accessibility or replication timing alone. Such loci may reflect the combined effects of positive selection and localized mutational processes not captured by the current epigenomic feature set, and therefore provide a focused starting point for downstream mechanistic and functional follow-up. Overall, integrative modeling of regional mutagenesis reveals both tissue-linked determinants of metastatic mutation landscapes and mutation-enriched loci beyond epigenomic expectation.

## Discussion

We integrated somatic mutation landscapes of metastatic cancers with tissue-resolved chromatin accessibility and replication timing profiles in CAMM to model regional mutation density across six cancer types. Two main points emerge from this analysis. First, metastatic genomes retain a strong tissue-of-origin epigenomic signature in their regional mutational landscape, with CA profiles from the tissue of origin consistently emerging as the most informative predictors, extending observations from primary cancer genomes ^22,23^. CA profiles from primary cancers remained informative despite being derived from independent cohorts rather than matched metastatic samples. This suggests that lineage-linked chromatin context remains a major determinant of regional mutagenesis even in advanced disease. Second, residuals from the epigenome-based model identify loci with mutation burden exceeding regional expectation, offering a complementary strategy to prioritize coding and non-coding genomic elements for further evaluation.

The persistence of tissue-matched CA as a dominant predictor in metastases suggests that key features of chromatin organization associated with cell identity remain stable during metastatic progression. One possible explanation is that epigenomic states are maintained through mitotic inheritance mechanisms, including histone modifications, DNA methylation, and chromatin remodeling, together with sustained activity of lineage-defining transcriptional programs ^23^. The directionality of the CA effect is also consistent with prior models of regional mutagenesis: more accessible regions tend to accumulate fewer mutations, whereas less accessible chromatin is associated with higher mutation burden, potentially reflecting differences in DNA damage and repair across chromatin states ^17,24,25^. Together, these observations are consistent with a continuing influence of chromatin context on metastatic regional mutagenesis.

Our findings extend prior work linking chromatin organization to regional mutation rates in primary tumors. Earlier studies showed that large-scale chromatin organization is associated with mutation rates at broad genomic scales, and that cell-of-origin CA shapes regional mutational landscapes across cancer types ^20,23^. The metastatic setting is not a trivial extension of these observations: metastatic tumors experience additional evolutionary selection during dissemination and are often exposed to therapies that leave distinct mutational footprints ^1,2,15,53^. Because the HMF cohort comprises treated metastatic cancers, therapy-associated mutagenesis may contribute variance that is not fully captured by CA or RT alone. Despite these influences, the strong tissue-matched CA contributions observed in metastatic genomes support the view that tissue-linked chromatin context remains a major determinant of regional mutation density even in advanced disease.

Methodologically, CAMM benefited from its multi-scale and multi-task architecture and consistently outperformed simpler baseline models. Its multi-scale design helps connect broad chromosomal domains to finer regional variability, which is difficult to capture in single-scale analyses ^13,14^. Whereas earlier epigenome-based models of regional mutagenesis generally analyzed mutation density at fixed genomic scales, CAMM jointly integrates megabase and kilobase-scale information, providing a richer view of how broad chromatin domains and local genomic context shape regional mutation density. Its transfer to independent primary tumors from PCAWG without retraining supports the broader applicability of the framework beyond the training cohort of metastatic cancers, although transfer to PCAWG is influenced by biological and technical differences between cohorts. CAMM also achieved useful indel prediction despite the lower frequency and greater sparsity of indels, although performance remained weaker than for SNVs, especially at finer resolution where local sequence determinants are less well captured by CA and RT features.

Beyond prediction, our residual-based framework provides a complementary way to prioritize mutation-enriched loci whose burden exceeds epigenome-based expectation. Existing genome-wide driver discovery approaches identify coding and non-coding regions with mutation frequencies above background expectation while accounting for covariates that shape background mutation rates ^9,54^. By explicitly modeling CA and RT as major determinants of regional mutagenesis ^13,14^, our approach highlights windows whose observed mutation density is not well explained by regional epigenomic context alone. The recovery of canonical cancer genes such as *TP53, PIK3CA*, or *APC* supports the biological relevance of this residual-based framework and its ability to recover known driver loci. The identification of loci such as the non-coding RNA *MALAT1* ^*52*^ and the oncogenic microRNA *MIR21* ^*50*^ suggests that the framework can also prioritize mutation-enriched non-coding regions with plausible cancer relevance. In this setting, residual outliers should be viewed as a prioritization signal for further analysis rather than as direct evidence of positive selection or driver status.

### Limitations of the study

This study has several limitations. First, although the models quantify strong associations between epigenomic features and regional mutation density, they do not establish causality. CA may influence mutation accumulation through DNA damage and repair processes, or both may reflect underlying transcriptional and replication-linked states that independently shape mutational patterns ^17,24^. Second, we relied on CA profiles from primary tumors rather than matched metastatic samples. Although these primary-tumor profiles remained strongly informative for metastatic mutation landscapes, matched metastatic epigenomes would likely improve biological specificity and help resolve metastasis-associated chromatin states more precisely. Third, the current framework considers only CA and RT, and therefore does not capture additional determinants of regional mutagenesis such as histone modifications, DNA methylation, 3D genome organization, or local sequence context, which may be particularly important at finer genomic scales ^10,11^. As these datasets become more widely available, integrating matched metastatic epigenomes and additional regulatory and sequence-level features should further improve both predictive performance and biological interpretation of regional mutagenesis.

### Conclusion

In summary, metastatic cancers retain a strong tissue-of-origin epigenomic signature in their regional mutation landscapes, indicating that chromatin context remains an important determinant of mutational heterogeneity in advanced disease. CAMM also offers a general framework for integrating epigenomic context into models of regional mutagenesis. Within this framework, residual analysis highlights coding and non-coding loci with mutation burden beyond epigenomic expectation. Together, these findings support an epigenome-informed view of metastatic cancer genomes.

## Methods

### Genomic datasets and quality control

We analyzed somatic mutations from the Hartwig Medical Foundation (HMF) cohort of metastatic solid tumors ^4^ and used the Pan-Cancer Analysis of Whole Genomes (PCAWG) cohort of primary tumors ^5^ for external validation. All sequencing and primary processing were performed by HMF under locally approved Institutional Review Board protocols with written informed consent, as described previously ^4^. This secondary analysis was approved by the University of Toronto Research Ethics Board (protocol 37521). For the HMF analysis, we used preprocessed whole-genome sequencing (WGS) variant calls aligned to GRCh37. We restricted analyses to autosomes, excluding sex-chromosome variants, and removed hypermutated tumors (>90,000 mutations). We focused on six metastatic cancer types with more than 150 samples (breast, colorectal, prostate, lung, esophagus, and skin), yielding 2,507 tumors comprising 45.4 million SNVs and 3.5 million indels. For validation in the PCAWG cohort, we selected six primary tumor cohorts matching the HMF dataset, applied the same autosome-only and hypermutation filtering, and all the cancer types have at least 50 samples. This resulted in 889 tumors with 9.3 million SNVs and 0.45 million indels. Both cohorts were analyzed in GRCh37 coordinates to ensure consistent genomic windows across the analyses.

### Epigenomic datasets

Chromatin accessibility (CA) profiles were obtained from prior TCGA studies performed using ATAC-seq experiments ^29^, comprising 796 profiles from 410 primary cancer samples and 404 unique patients. We specifically selected cancer-derived profiles, excluding normal tissue samples to maintain relevance to tumor mutational patterns. ATAC-seq peak coordinates were converted from GRCh38 to GRCh37 using the UCSC LiftOver software ^55^ with default chain files. Replication timing (RT) data representing 96 cell lines were retrieved from the ENCODE RepliSeq project ^30^ (GEO: GSE34399). RT profiles were generated by calculating mean signal intensities across genomic windows, with early and late replication domains defined by signal quantiles. The combined epigenomic dataset comprised 892 profiles in total, including 796 CA and 96 RT profiles.

### Genomic windows and feature construction

To integrate mutation density with epigenomic context, we partitioned the autosomal genome into non-overlapping windows at 10 kb, 100 kb, and 1 Mb resolutions. Larger windows (100 kb, 1 Mb) were constructed by aggregating consecutive 10 kb windows. We excluded windows with low mappability (<80%, based on UMAP mappability tracks) ^56^, windows overlapping ENCODE blacklist regions ^57^, and windows on sex chromosomes. After filtering, we retained 264,105 windows at 10 kb, 26,401 windows at 100 kb, and 2,629 windows at 1 Mb resolution. Next, for each epigenomic profile and each genomic window, we computed the mean CA or RT signals within that window, producing a window-by-feature matrix at each resolution. For the response variables, we quantified regional mutational density separately for each cancer type by counting the total number of SNVs and indels within each window across all tumors of that cancer type. This window-level formulation yields stable estimates of regional mutation density and defines the unit of prediction used throughout model training and evaluation.

### Hierarchical multi-scale, multi-task model architecture

We developed CAMM, a hierarchical multi-scale and multi-task neural network that jointly predicts SNV and indel burdens at three genomic resolutions (1 Mb, 100 kb, and 10 kb). The model has three components. First, for scale-specific feature extraction, a CA/RT feature vector at each resolution is passed through an adaptive gating layer ^58^ (linear layer plus sigmoid) to learn element-wise feature weights. The gated features are then processed by a four-layer multilayer perceptron (MLP) with ReLU activations, batch normalization ^59^, and dropout ^60^ to generate a scale-specific embedding. Second, to enable coarse-to-fine information flow across scales, coarser-scale embeddings are propagated to finer resolutions to capture hierarchical dependence of mutation rates across genomic scales ^31^. A shared fully connected block transforms the 1 Mb embedding into a latent representation, *h*_*1*_, which is used both for 1 Mb prediction and as contextual input to finer scales. At 100 kb, the corresponding embedding is concatenated with h_1_ and transformed into h_2_, which supports 100 kb prediction and provides additional context for 10 kb. At 10 kb, the 10 kb embedding is concatenated with *h*_*1*_ and *h*_*2*_, and then passed into two task-specific towers to produce *h*_*3*_^*SNV*^ and *h*_*3*_^*INDEL*^ representations. Third, for prediction heads and multi-task weighting, we attached task- and scale-specific linear prediction heads for SNV and indel prediction at each resolution. Outputs were passed through a Softplus activation to enforce non-negative predictions. Losses across tasks and scales were combined using uncertainty-based weighting ^61^, with learnable task-specific uncertainty parameters that adaptively weight SNV versus indel objectives during optimization, mitigating domination of the objective by the more frequent SNV task ^62^.

### Model training, cross-validation, and evaluation

Models were trained separately for each cancer type, with performance estimated using 5-fold cross-validation repeated 12 times; within each run, 80% of windows were used for training and 20% for validation. The same repeated cross-validation splits were used when comparing the full multi-task, multi-scale model against single-scale and single-task models, random forest baselines, and elastic net baselines. Error bars in figures represent the standard deviation across cross-validation folds. Hyperparameter optimization was performed using Optuna ^63^ with 50 trials per cancer type, optimizing the learning rate (10^-5^ to 10^-2^), batch size (32 to 256), hidden dimensions (64 to 512), and dropout rate (0.1 to 0.5). Training employed the Adam optimizer ^64^ with early stopping (patience = 10 epochs) based on validation loss. To further assess robustness to genomic autocorrelation among neighboring windows, we performed chromosome hold-out validation. For each autosome (chromosomes 1-22), we trained the model using windows from the other 21 autosomes and evaluated performance on the held-out chromosome.

### External validation on PCAWG primary tumors

To assess cross-cohort generalizability, we applied CAMM trained on HMF ^4^ to an independent set of primary tumor genomes from the PCAWG project ^5^. Because absolute mutation burdens can differ between metastatic and primary cohorts owing to biological and technical factors, we evaluated external validation using non-parametric Spearman correlation between observed and predicted regional mutation density across windows, reported by cancer type and scale.

### Feature importance and model interpretation

We assessed epigenomic feature importance using a combination of permutation testing and Shapley-based attribution. For permutation-based feature importance, we computed the decrease in predictive performance after permuting each CA or RT feature across held-out validation windows (ΔR^2^ = R^2^baseline - R^2^permuted). Permutations were repeated 1,000 times per feature to obtain a null distribution of ΔR^2^ values. Empirical one-sided P-values were computed with add-one smoothing:

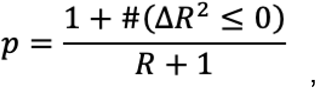

where smaller p-values indicate important features whose permutation consistently reduces performance. Features with empirical P < 0.001 were considered significant. Next, to characterize the direction and magnitude of each feature’s contribution to individual regional predictions, we computed Shapley Additive Explanation (SHAP) ^35^ values using Captum’s ShapleyValueSampling implementation ^65^. Features were ranked by mean absolute SHAP value, and top features were defined as SHAP top-30 predictors that also reached permutation significance (P < 0.001). We additionally examined correlations between feature values and their SHAP attributions to summarize whether higher CA or RT values tended to increase or decrease predicted mutation density. To interpret feature importance in the context of cell of origin and formally test enrichment of tissue-linked CA profiles among top predictors, each CA profile was classified for each cancer type as tissue-matched, anatomically related, or unrelated. The full set of 796 CA profiles was used as the background feature universe, while RT profiles were excluded from CA enrichment analyses. We tested enrichment of tissue-matched and tissue-matched-or-related CA profiles among selected predictors using one-sided Fisher’s exact tests. To compare attribution magnitudes across the full CA feature set, we used one-sided Wilcoxon rank-sum tests to assess whether tissue-matched or tissue-matched-or-related CA profiles had larger mean absolute SHAP values than unrelated profiles. P-values were adjusted using the Benjamini-Hochberg procedure.

### Identification of mutation-enriched windows and cancer gene annotation

To identify regions where observed mutation burden exceeded epigenome-based expectations, we first log-transformed CAMM residuals between observed and predicted mutation counts for each 10 kb window. Standardized residuals (Z-scores) were then computed from the transformed residual distribution. To assess the effect of threshold choice, we tested multiple Z-score cutoffs, including Z > 2, Z > 3, Z > 4, and Z > 5. We selected Z > 4 as the primary cutoff for downstream analysis because it provided a stringent definition of mutation-enriched windows while retaining enough loci for gene-level annotation. Windows with Z > 4 were therefore classified as mutation-enriched, indicating that more mutations were observed in these windows than predicted by the epigenome-based model, as reflected by positive residuals. Mutation-enriched windows were annotated using GRCh37 gene coordinates from NCBI GFF annotations. We assigned windows to genes based on genomic overlap and compiled the resulting gene list. To prioritize cancer-relevant loci, we cross-referenced genes against the COSMIC Cancer Gene Census ^40^ and OncoKB ^41^ databases, and retained genes present in at least one database for downstream reporting.

## Supplementary tables

**Table S1. Summary of metastatic tumor mutation data from the Hartwig Medical Foundation (HMF) cohort**. This table shows the number of tumor samples and total mutation counts for each cancer type included in the study.

Single-nucleotide variants (SNVs) and insertions/deletions (indels) are reported separately for breast, colorectal, esophagus, lung, prostate, and skin cancers, with totals across all cancer types.

**Table S2. Summary of chromatin accessibility (CA) and DNA replication timing (RT) profiles used in the study**. This table lists the CA and RT predictors used for modeling. For each predictor, the table provides the assay or technique, predictor name, sample or profile description, predictor category, related the cancer type from HMF where applicable, and the original study source.

**Table S3. Summary of primary tumor mutation data from the Pan-Cancer Analysis of Whole Genomes (PCAWG) project**. This table shows the number of tumor samples and total mutation counts for each cancer type included in the primary tumor cohort. SNVs and indels are reported separately for breast, colorectal, esophagus, lung, prostate, and skin cancers, with totals across all cancer types.

**Table S4. Summary of outlier genomic windows identified by the CAMM model**. This table lists genomic windows with elevated mutation counts identified by CAMM across cancer types and variant classes. For each window, the table provides genomic coordinates, window size, observed and predicted mutation counts, residual-based enrichment statistics, gene overlap information, overlapping gene annotations, and whether the window contains a known cancer gene.

## Data availability

Processed PCAWG mutation data and chromatin accessibility/replication timing (CA/RT) feature matrices used in this study are available in our GitHub repository: https://github.com/reimandlab/CAMM. Input WGS data and metadata annotations for metastatic cancer samples from the Hartwig Medical Foundation (HMF) are controlled-access datasets. Access to these data can be requested from the HMF and are subject to scientific review and completion of the required data access or material transfer agreements. Intermediate files derived from HMF controlled-access datasets are not publicly shared because of data-use restrictions. All publicly shareable processed data required to reproduce the main analyses are provided in the repository where permitted by the original data-use terms.

## Code availability

All code used for data preprocessing, model training, downstream analysis, and visualization is available in our GitHub repository: https://github.com/reimandlab/CAMM. Documentation for reproducing the main analyses and figures is also included

## Acknowledgments

We would like to thank Zoe P. Klein and Jigyansa Mishra for valuable discussions. This work was partially supported by the Canadian Institutes of Health Research (CIHR) Project Grant (PJT-197925) to J.R. and the Investigator Award to J.R. from the Ontario Institute for Cancer Research (OICR). H.J. was partially supported by the Computer Science Engagement Award from the Department of Computer Science, University of Toronto and the Undergraduate Research Fund from the Faculty of Arts and Science, University of Toronto. A.B. was partially supported by the Ontario Graduate Scholarship (OGS). K.C. was partially supported by fellowships from the Medical Biophysics Department at University of Toronto. Funding to OICR is provided by the Government of Ontario. This publication and the underlying study have been made possible partly based on the data that Hartwig Medical Foundation has made available to the study.

## Author Contributions

H.J. developed the computational model, analysed the data, and prepared the figures with input from J.R.. H.J. and J.R. wrote the manuscript. A.T.B. and K.C.L.C. contributed to data analyses and interpretation. J.R. conceptualized the idea, supervised the project, and acquired funding. All coauthors reviewed, edited and approved the final manuscript.

## References

1. Lambert, A. W., Pattabiraman, D. R. & Weinberg, R. A. Emerging biological principles of metastasis. Cell 168, 670–691 (2017).

2. Massagué, J. & Obenauf, A. C. Metastatic colonization by circulating tumour cells. Nature 529, 298–306 (2016).

3. Welch, D. R. & Hurst, D. R. Defining the hallmarks of metastasis. Cancer Res. 79, 3011–3027 (2019).

4. Priestley, P. et al. Pan-cancer whole-genome analyses of metastatic solid tumours. Nature 575, 210–216 (2019).

5. ICGC/TCGA Pan-Cancer Analysis of Whole Genomes Consortium. Pan-cancer analysis of whole genomes. Nature 578, 82–93 (2020).

6. Gerstung, M. et al. The evolutionary history of 2,658 cancers. Nature 578, 122–128 (2020).

7. Martincorena, I. et al. Universal patterns of selection in cancer and somatic tissues. Cell 171, 1029–1041.e21 (2017).

8. Rheinbay, E. et al. Analyses of non-coding somatic drivers in 2,658 cancer whole genomes. Nature 578, 102–111 (2020).

9. Zhu, H. et al. Candidate cancer driver mutations in distal regulatory elements and long-range chromatin interaction networks. Mol. Cell 77, 1307–1321.e10 (2020).

10. Alexandrov, L. B. et al. The repertoire of mutational signatures in human cancer. Nature 578, 94–101 (2020).

11. Li, Y. et al. Patterns of somatic structural variation in human cancer genomes. Nature 578, 112–121 (2020).

12. Kumar, S. et al. Passenger mutations in more than 2,500 cancer genomes: Overall molecular functional impact and consequences. Cell 180, 915–927.e16 (2020).

13. Supek, F. & Lehner, B. Scales and mechanisms of somatic mutation rate variation across the human genome. DNA Repair (Amst.) 81, 102647 (2019).

14. Gonzalez-Perez, A., Sabarinathan, R. & Lopez-Bigas, N. Local determinants of the mutational landscape of the human genome. Cell 177, 101–114 (2019).

15. Pich, O. et al. The mutational footprints of cancer therapies. Nat. Genet. 51, 1732–1740 (2019).

16. Katainen, R. et al. CTCF/cohesin-binding sites are frequently mutated in cancer. Nat. Genet. 47, 818–821 (2015).

17. Sabarinathan, R., Mularoni, L., Deu-Pons, J., Gonzalez-Perez, A. & López-Bigas, N. Nucleotide excision repair is impaired by binding of transcription factors to DNA. Nature 532, 264–267 (2016).

18. Uusküla-Reimand, L. et al. Topoisomerase IIb binding delineates localized mutational processes and driver mutations in cancer genomes. Nat. Commun. 16, 10241 (2025).

19. Lawrence, M. S. et al. Mutational heterogeneity in cancer and the search for new cancer-associated genes. Nature 499, 214–218 (2013).

20. Schuster-Böckler, B. & Lehner, B. Chromatin organization is a major influence on regional mutation rates in human cancer cells. Nature 488, 504–507 (2012).

21. Stamatoyannopoulos, J. A. et al. Human mutation rate associated with DNA replication timing. Nat. Genet. 41, 393–395 (2009).

22. Ocsenas, O. & Reimand, J. Chromatin accessibility of primary human cancers ties regional mutational processes and signatures with tissues of origin. PLoS Comput. Biol. 18, e1010393 (2022).

23. Polak, P. et al. Cell-of-origin chromatin organization shapes the mutational landscape of cancer. Nature 518, 360–364 (2015).

24. Supek, F. & Lehner, B. Differential DNA mismatch repair underlies mutation rate variation across the human genome. Nature 521, 81–84 (2015).

25. Zheng, C. L. et al. Transcription restores DNA repair to heterochromatin, determining regional mutation rates in cancer genomes. Cell Rep. 9, 1228–1234 (2014).

26. Woo, Y. H. & Li, W.-H. DNA replication timing and selection shape the landscape of nucleotide variation in cancer genomes. Nat. Commun. 3, 1004 (2012).

27. Liu, L., De, S. & Michor, F. DNA replication timing and higher-order nuclear organization determine single-nucleotide substitution patterns in cancer genomes. Nat. Commun. 4, 1502 (2013).

28. Jiao, W. et al. A deep learning system accurately classifies primary and metastatic cancers using passenger mutation patterns. Nat. Commun. 11, 728 (2020).

29. Corces, M. R. et al. The chromatin accessibility landscape of primary human cancers. Science 362, eaav1898 (2018).

30. ENCODE Project Consortium et al. Expanded encyclopaedias of DNA elements in the human and mouse genomes. Nature 583, 699–710 (2020).

31. Ronneberger, O., Fischer, P. & Brox, T. U-Net: Convolutional Networks for Biomedical Image Segmentation. in Lecture Notes in Computer Science 234–241 (Springer International Publishing, Cham, 2015). doi:10.1007/978-3-319-24574-4_28.

32. Robinson, D. et al. Integrative clinical genomics of advanced prostate cancer. Cell 161, 1215–1228 (2015).

33. Rebello, R. J. et al. Prostate cancer. Nat. Rev. Dis. Primers 7, 9 (2021).

34. Martínez-Jiménez, F. et al. Pan-cancer whole-genome comparison of primary and metastatic solid tumours. Nature 618, 333–341 (2023).

35. Lundberg, S. & Lee, S.-I. A unified approach to interpreting model predictions. arXiv [cs.AI] (2017).

36. Rhind, N. & Gilbert, D. M. DNA replication timing. Cold Spring Harb. Perspect. Biol. 5, a010132 (2013).

37. Hiratani, I. et al. Global reorganization of replication domains during embryonic stem cell differentiation. PLoS Biol. 6, e245 (2008).

38. Ryba, T. et al. Evolutionarily conserved replication timing profiles predict long-range chromatin interactions and distinguish closely related cell types. Genome Res. 20, 761–770 (2010).

39. Sherman, M. A. et al. Genome-wide mapping of somatic mutation rates uncovers drivers of cancer. Nat. Biotechnol. 40, 1634–1643 (2022).

40. Sondka, Z. et al. The COSMIC Cancer Gene Census: describing genetic dysfunction across all human cancers. Nat. Rev. Cancer 18, 696–705 (2018).

41. Chakravarty, D. et al. OncoKB: A precision oncology knowledge base. JCO Precis. Oncol. 2017, (2017).

42. Samuels, Y. et al. High frequency of mutations of the PIK3CA gene in human cancers. Science 304, 554 (2004).

43. Grinshpun, A., Chen, V., Sandusky, Z. M., Fanning, S. W. & Jeselsohn, R. ESR1 activating mutations: From structure to clinical application. Biochim. Biophys. Acta Rev. Cancer 1878, 188830 (2023).

44. Chen, X. et al. XBP1 promotes triple-negative breast cancer by controlling the HIF1α pathway. Nature 508, 103–107 (2014).

45. Fearon, E. R. & Vogelstein, B. A genetic model for colorectal tumorigenesis. Cell 61, 759–767 (1990).

46. Cancer Genome Atlas Network. Comprehensive molecular characterization of human colon and rectal cancer. Nature 487, 330–337 (2012).

47. Campos, E. I., Martinka, M., Mitchell, D. L., Dai, D. L. & Li, G. Mutations of the ING1 tumor suppressor gene detected in human melanoma abrogate nucleotide excision repair. Int. J. Oncol. 25, 73–80 (2004).

48. Kalashnikova, E. V. et al. ANCCA/ATAD2 overexpression identifies breast cancer patients with poor prognosis, acting to drive proliferation and survival of triple-negative cells through control of B-Myb and EZH2. Cancer Res. 70, 9402–9412 (2010).

49. Ciró, M. et al. ATAD2 is a novel cofactor for MYC, overexpressed and amplified in aggressive tumors. Cancer Res. 69, 8491–8498 (2009).

50. Bautista-Sánchez, D. et al. The promising role of miR-21 as a cancer biomarker and its importance in RNA-based therapeutics. Mol. Ther. Nucleic Acids 20, 409–420 (2020).

51. Wang, C. et al. An autoregulatory feedback loop of miR-21/VMP1 is responsible for the abnormal expression of miR-21 in colorectal cancer cells. Cell Death Dis. 11, 1067 (2020).

52. Kim, J. et al. Long noncoding RNA MALAT1 suppresses breast cancer metastasis. Nat. Genet. 50, 1705–1715 (2018).

53. Cheng, K. C. L. et al. Therapy-associated mutagenesis at CTCF binding sites is shaped by chromatin context and DNA repair capacity. bioRxivorg (2026) doi:10.64898/2026.04.15.718780.

54. Shuai, S., PCAWG Drivers and Functional Interpretation Working Group, Gallinger, S., Stein, L. D. & PCAWG Consortium. Combined burden and functional impact tests for cancer driver discovery using DriverPower. Nat. Commun. 11, 734 (2020).

55. Kuhn, R. M., Haussler, D. & Kent, W. J. The UCSC genome browser and associated tools. Brief. Bioinform. 14, 144–161 (2013).

56. Karimzadeh, M., Ernst, C., Kundaje, A. & Hoffman, M. M. Umap and Bismap: quantifying genome and methylome mappability. Nucleic Acids Res. 46, e120 (2018).

57. Amemiya, H. M., Kundaje, A. & Boyle, A. P. The ENCODE blacklist: Identification of problematic regions of the genome. Sci. Rep. 9, 9354 (2019).

58. Dauphin, Y. N., Fan, A., Auli, M. & Grangier, D. Language modeling with gated convolutional networks. arXiv [cs.CL] (2016).

59. Ioffe, S. & Szegedy, C. Batch Normalization: Accelerating deep network training by reducing internal covariate shift. arXiv [cs.LG] (2015).

60. Srivastava, N., Hinton, G. E., Krizhevsky, A., Sutskever, I. & Salakhutdinov, R. Dropout: a simple way to prevent neural networks from overfitting. J. Mach. Learn. Res. 15, 1929–1958 (2014).

61. Cipolla, R., Gal, Y. & Kendall, A. Multi-task learning using uncertainty to weigh losses for scene geometry and semantics. in 2018 IEEE/CVF Conference on Computer Vision and Pattern Recognition (IEEE, 2018). doi:10.1109/cvpr.2018.00781.

62. Martincorena, I. & Campbell, P. J. Somatic mutation in cancer and normal cells. Science 349, 1483–1489 (2015).

63. Akiba, T., Sano, S., Yanase, T., Ohta, T. & Koyama, M. Optuna: A next-generation hyperparameter optimization framework. arXiv [cs.LG] (2019).

64. Kingma, D. P. & Ba, J. Adam: A method for stochastic optimization. arXiv [cs.LG] (2014).

65. Kokhlikyan, N. et al. Captum: A unified and generic model interpretability library for PyTorch. arXiv [cs.LG] (2020).

