## Supplementary Figures for "Chromatin accessibility of primary cancers informs regional mutagenesis in metastases through multi-scale deep learning"

### Supplementary materials

#### Jiang et al. (2026)

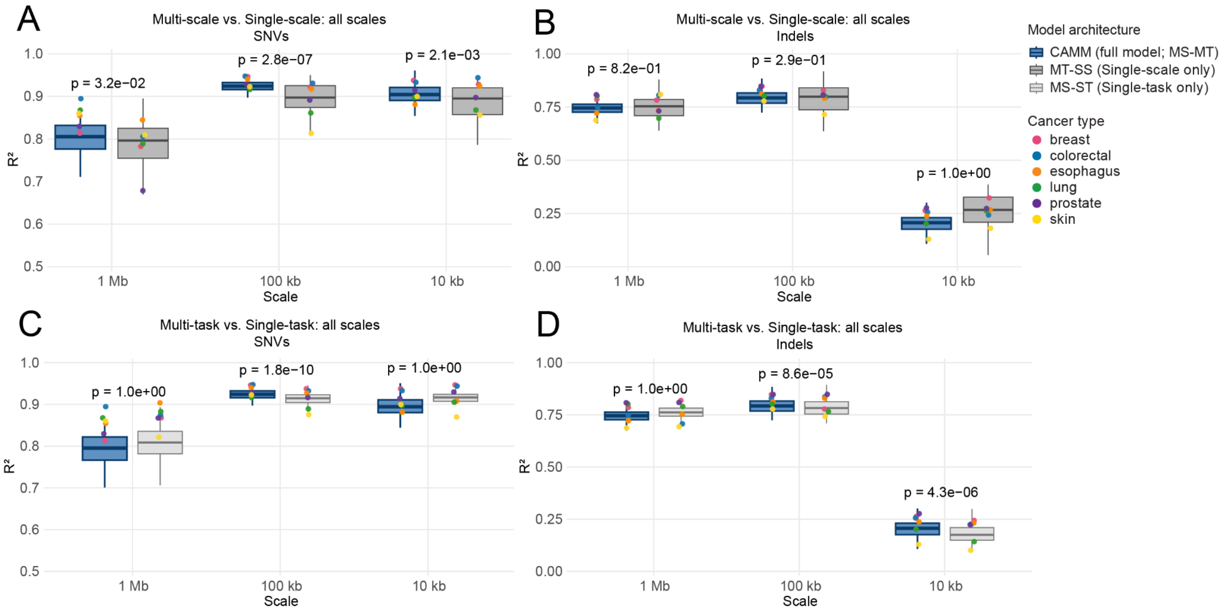


**Figure S1. Benchmarking CAMM against single-scale and single-task baselines for epigenome-based prediction of regional mutation density across genomic scales. (A-B)** Performance of the multi-scale, multi-task model CAMM (MS-MT) compared with the multi-task, single-scale model (MT-SS) for predicting regional mutation density from epigenomic features across 1 Mb, 100 kb, and 10 kb genomic bins for **(A)** single-nucleotide variants (SNVs) and **(B)** insertions and deletions (indels). **(C-D)** Performance of CAMM compared with the multi-scale, single-task model (MS-ST) across the same genomic scales for **(C)** SNVs and **(D)** indels. Prediction performance is reported as the coefficient of determination ($R^2$) between predicted and observed mutation density. Boxplots summarize performance across cancer types; colored points denote individual cancer types: breast, colorectal, esophagus, lung, prostate, and skin. P-values above each scale indicate one-sided statistical comparisons testing whether the MS-MT model outperformed the corresponding baseline model.

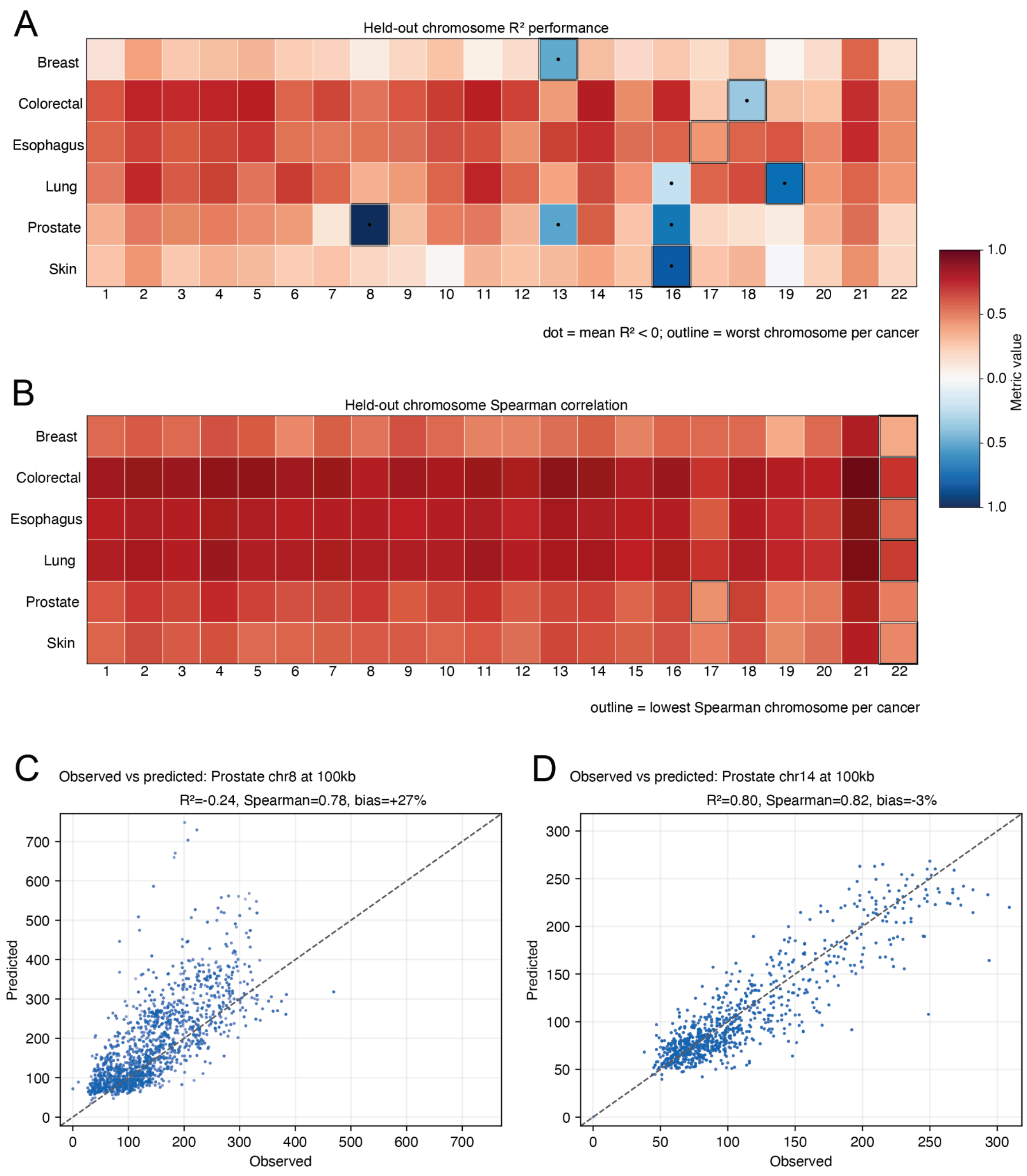


**Figure S2. Chromosome-held-out validation of CAMM assessing model robustness under chromosome-level holdout rather than random window splitting. (A)** Heatmap of chromosome-held-out performance ($R^2$) across cancer types. For each cancer type and held-out autosome, $R^2$ values were averaged across genomic resolutions and mutation types. Each column represents an autosome, and each row represents a cancer type. Dots indicate chromosomes with mean $R^2 < 0$, and outlined cells mark the worst-performing chromosome for each cancer type. **(B)** Corresponding chromosome-held-out Spearman correlations, averaged across genomic resolutions and mutation types; outlined cells indicate the lowest-correlation chromosome for each cancer type. **(C-D)** Representative examples from analyses of prostate cancer genomes comparing observed and predicted SNV burden across 100-kb windows for **(C)** chromosome 8, showing poor calibration but preserved rank correlation, and **(D)** chromosome 14, showing strong agreement of observed and predicted values. Dashed lines indicate the identity line.

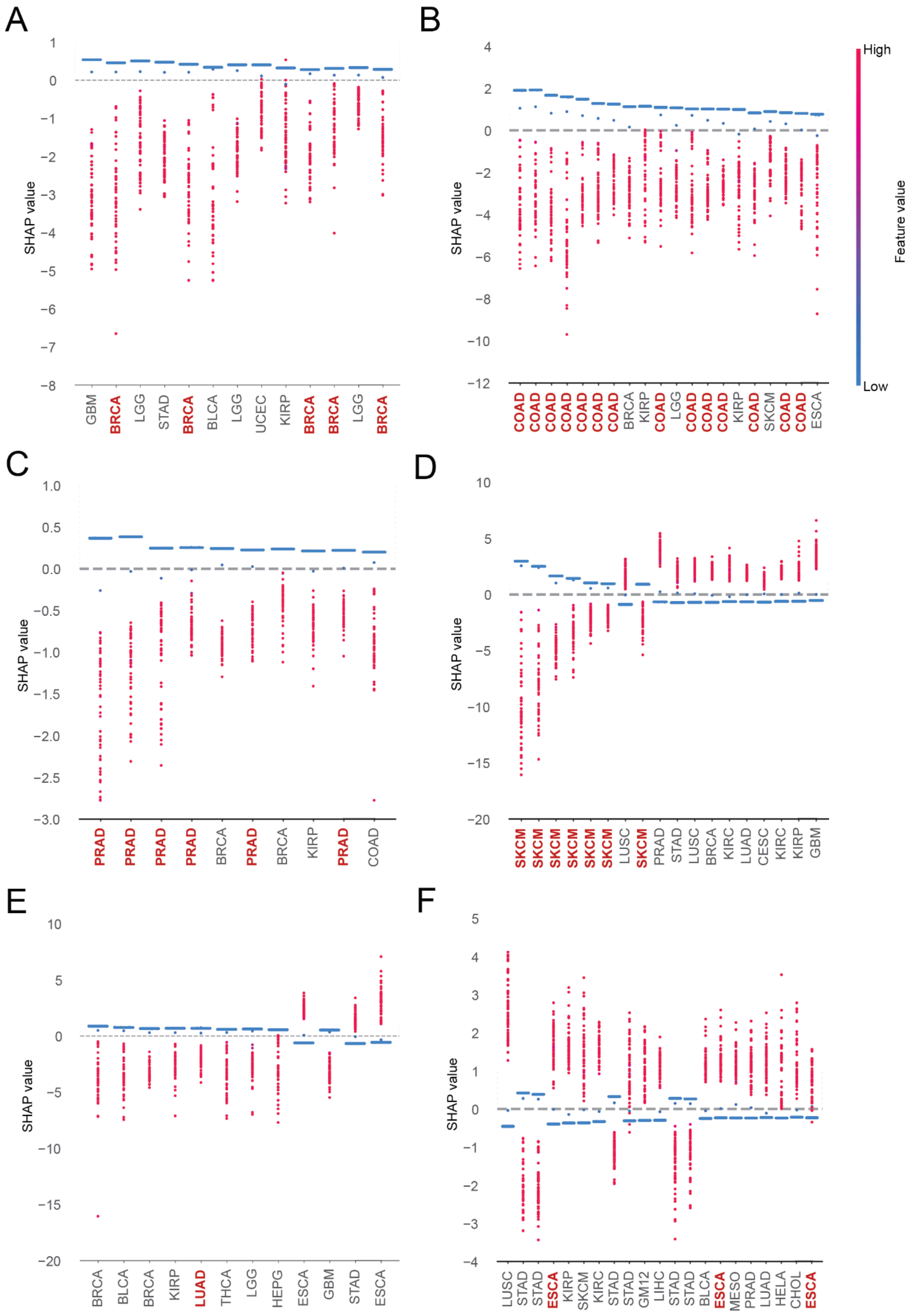
A
SHAP value
GBM BRCA LGG STAD BRCA BLCA LGG UCEC KIRP BRCA BRCA LGG BRCA
B
SHAP value
COAD COAD COAD COAD COAD COAD COAD BRCA KIRP COAD LGG COAD COAD COAD KIRP COAD SKCM COAD COAD ESCA
High
Feature value
Low
C
SHAP value
PRAD PRAD PRAD PRAD BRCA PRAD BRCA KIRP PRAD COAD
D
SHAP value
SKCM SKCM SKCM SKCM SKCM SKCM LUSC SKCM PRAD STAD LUSC BRCA KIRC LUAD CESC KIRC KIRP GBM
E
SHAP value
BRCA BLCA BRCA KIRP LUAD THCA LGG HEPG ESCA GBM STAD ESCA
F
SHAP value
LUSC STAD STAD ESCA KIRP SKCM KIRC STAD STAD GM12 LIHC STAD STAD BLCA ESCA MESO PRAD LUAD HELA CHOL ESCA

**Figure S3. SHAP-based interpretation of significant CA and RT predictors in regional mutagenesis. (A-F)** SHAP attribution scores for significant chromatin accessibility (CA) and replication timing (RT) predictors in **(A)** breast, **(B)** colorectal, **(C)** prostate, **(D)** skin, **(E)** lung, and **(F)** esophageal metastases. Predictors were selected by permutation-based importance testing and ranked by mean absolute SHAP value.

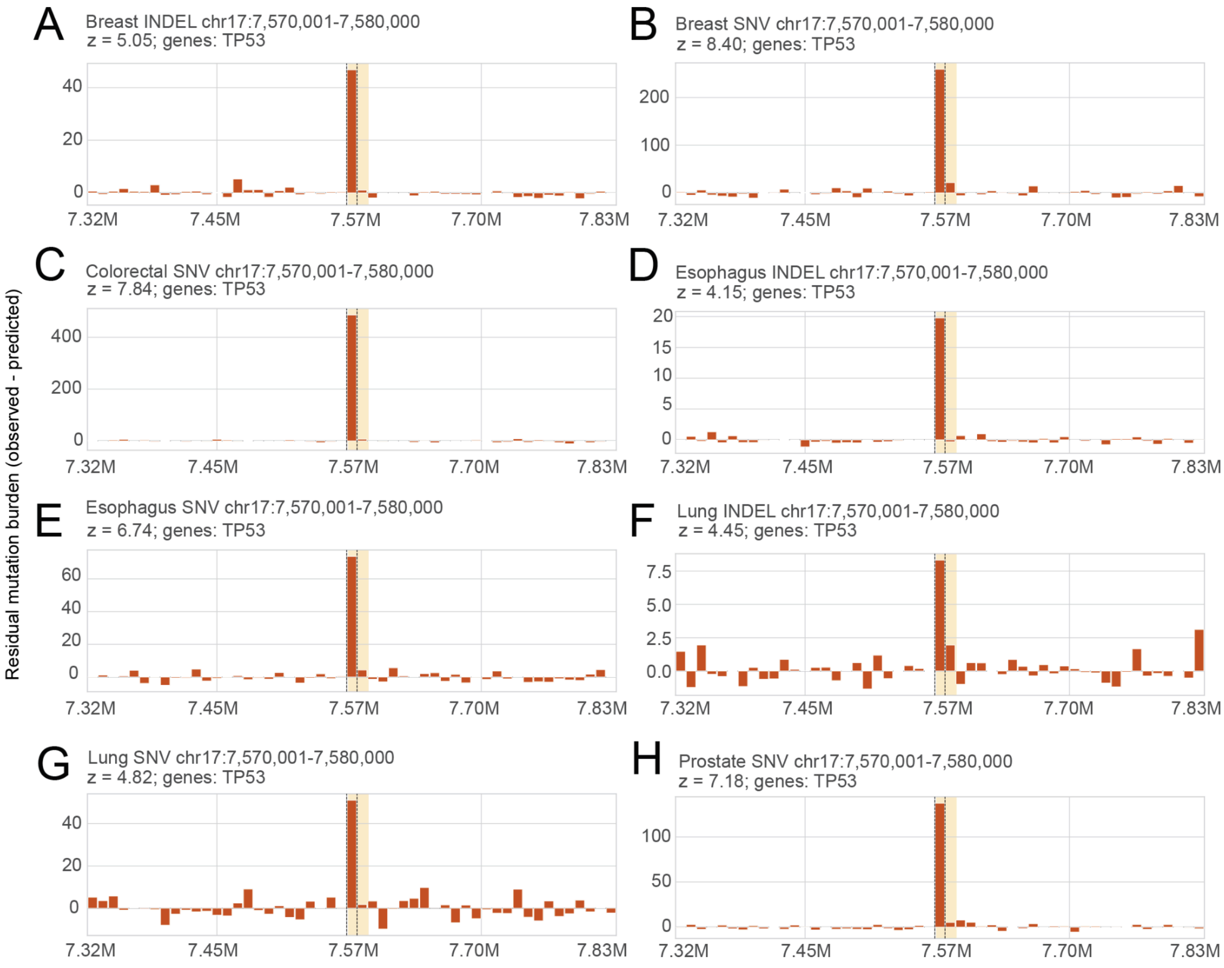


**Figure S4. Mutation-enriched genomic windows at the *TP53* locus. (A-H)** Residual profiles surrounding the TP53 locus at 10-kb resolution for combinations of cancer types and mutation classes in which the genomic window overlapping TP53 was identified as a high-positive-residual outlier. Bars show model residuals, calculated as observed minus predicted mutation density, across neighboring 10-kb genomic windows. The highlighted region marks the 10-kb window overlapping *TP53*. Positive residuals indicate windows in which observed mutation density exceeded epigenome-based predictions, consistent with mutation enrichment beyond the expected CA and RT background.

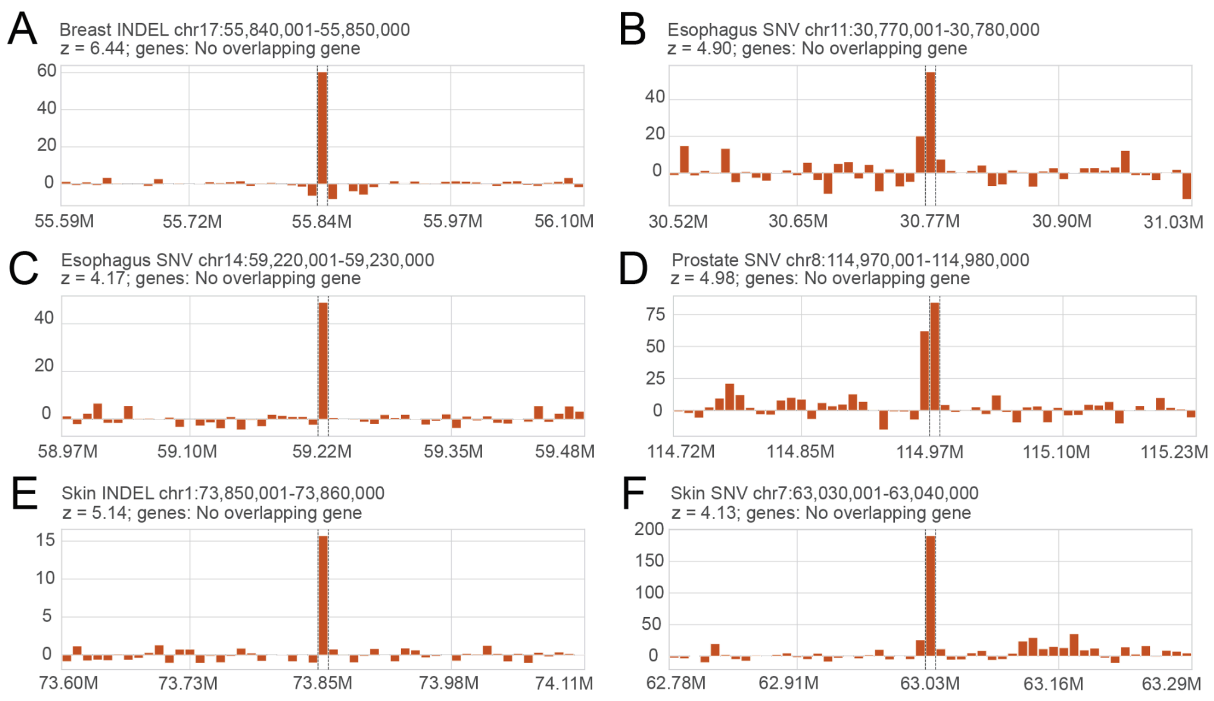


**Figure S5.** Examples of underestimated 10-kb windows with no overlapping genes. **(A-F)** Representative residual profiles surrounding high-positive-residual 10 kb windows that did not overlap annotated genes.
